# Phagocytosis deficient glia display phagosome processing defects and macrophage recruitment to the brain of adult *Drosophila melanogaster*

**DOI:** 10.64898/2026.02.20.707075

**Authors:** Guangmei Liu, Iqra Amin, Cheng Yang Shi, Kimberly McCall

**Affiliations:** Department of Biology, Boston University, 5 Cummington Mall, Boston, MA 02215, USA

## Abstract

Efficient clearance of dying cells is essential for brain homeostasis, yet how partial defects in phagocytic processing affect neuroimmune interactions during aging remains unclear. In the adult *Drosophila* brain, glia function as professional phagocytes through the conserved engulfment receptor Draper (Drpr). Here, we show that glial loss of Drpr does not completely eliminate phagocytosis but instead leads to persistent, age-dependent inefficiency in corpse degradation. Using a genetically encoded pH-sensitive reporter to visualize acidified phagocytic compartments, we find that *drpr*-deficient glia retain residual engulfment activity but progressively accumulate enlarged, incompletely degraded phagocytic cargo. This chronic clearance defect coincides with altered immune dynamics at the central nervous system periphery, including increased recruitment and adhesion of peripheral hemocytes at the blood–brain barrier (BBB), without overt BBB disruption. Notably, hemocytes at the brain surface can phagocytose glial material in a Drpr-dependent manner, revealing a form of barrier-associated “border clearance”. Together, these findings demonstrate that inefficient corpse degradation is sufficient to reshape neuroimmune interactions during aging.

**Main points:**

- Draper-deficient glia still engulf corpses but fail phagolysososomal degradation.
- Impaired glial clearance reshapes immune dynamics at the CNS periphery.
- With intact BBB, hemocytes that accumulate on the aging brain surface engulf glial debris.

## Introduction

Efficient clearance of dying cells is essential for maintaining tissue homeostasis. In the central nervous system (CNS) of both vertebrates and invertebrates, glial cells serve as professional phagocytes, continuously pruning synapses, removing apoptotic neurons, and clearing degenerating axons (Awasaki et al., 2006; Bishop et al., 2004; Chung et al., 2013; Galloway et al., 2019; Iram et al., 2016; Marín-Teva et al., 2004; Melcarne et al., 2019; Schafer et al., 2012). Phagocytosis is a multistep process that includes sensing of dying cells, receptor-mediated recognition, internalization, and lysosome-dependent degradation, with each stage regulated by distinct signaling pathways. Dying cells release chemotactic “find-me” signals and expose “eat me” cues on the surface, such as phosphatidylserine (PS) to promote engulfment by phagocytes (Appelt et al., 2005; Botelho & Grinstein, 2011; Segawa & Nagata, 2015). Recognition of these cues triggers cytoskeletal rearrangements that drive formation of phagosomes (Melcarne et al., 2019), which subsequently mature through sequential fusion with endosomes and lysosomes to form acidic phagolysosomes that degrade engulfed material (Botelho & Grinstein, 2011; Melcarne et al., 2019). Disruption of any step in this pathway compromises neural circuit integrity and is increasingly associated with chronic neuroinflammation and neurodegenerative diseases (Elguero et al., 2023; Galloway et al., 2019; Grau et al., 2015; Ni et al., 2024; Parhizkar et al., 2019; Tremblay et al., 2019). Despite these connections, most studies have focused on conditions in which phagocytosis is completely blocked, leaving unresolved how partial or progressive defects in corpse processing influence CNS homeostasis and neuroimmune interactions during aging.

*Drosophila* provides a powerful model to address this problem, as core mechanisms governing phagocytosis, lysosomal degradation and innate immune signaling are highly conserved, and aging-associated immune changes can be examined in vivo with cellular resolution (Melcarne et al., 2019). In the fly CNS, the engulfment receptor Draper (Drpr), an ortholog of mammalian MEGF10, serves as a central mediator of glial phagocytosis (Doherty et al., 2009; Logan et al., 2012; MacDonald et al., 2006). In mammals, MEGF10 is enriched in astrocytes and contributes to engulfment of synapses and apoptotic neurons in both developing and adult CNS (Chung et al., 2013; Iram et al., 2016). In *Drosophila*, Drpr is required for clearance of apoptotic neurons during development, removal of axonal debris after injury, and maintenance of adult brain homeostasis (Doherty et al., 2014; Freeman et al., 2003; Logan et al., 2012; Lu et al., 2017; MacDonald et al., 2006). Loss of *drpr* in the null allele *drpr*^Δ*5*^ disrupts phagosome-lysosome fusion and impairs phagosomal acidification and maturation in hemocytes (Petrignani et al., 2021) and glia (Etchegaray et al., 2016; Shklover et al., 2015), indicating a role in phagocytic processing rather than cargo recognition alone. Consistent with this role, activation of the JNK pathway downstream of Draper in embryonic glia restores phagosome number and enhances phagocytic activity (Shklover et al., 2015). Glial *drpr* knockdown leads to accumulation of cellular corpses and upregulation of innate immune signaling within the brain (Elguero et al., 2023; Etchegaray et al., 2016). However, the number of TUNEL-positive apoptotic cells in *drpr*-deficient brains does not increase indefinitely with age (Etchegaray et al., 2016), suggesting that apoptotic cell production and clearance may reach a dynamic equilibrium even when canonical engulfment signaling is compromised. Whether this equilibrium reflects persistent but inefficient phagocytic activity, or compensation by alternative cell types or clearance pathways, has remained unresolved.

This study tests whether partial defects in glial phagocytosis are sufficient to alter neuroimmune interactions during aging. In aging and disease, compromised clearance of dying cells and proteinaceous debris within the CNS is frequently linked to heightened neuroimmune crosstalk, including the recruitment of peripheral immune cells to neural tissues and disruption of blood-brain barrier (BBB) integrity, both of which can exacerbate inflammation and cellular stress (Kelly et al., 2013; Minogue et al., 2014; Stalder et al., 2005; Togo et al., 2002; Wyss-Coray & Rogers, 2012). Resolving how partial defects in resident phagocytic clearance influence immune dynamics during aging is therefore critical for understanding how CNS homeostasis erodes over time.

Across both mammals and invertebrates, disruptions in tissue homeostasis are often accompanied by recruitment of peripheral immune cells, raising the possibility that reduced efficiency of resident phagocytes may signal the need for immune compensation. In *Drosophila*, circulating hemocytes serve as professional phagocytes in peripheral tissues. Hemocytes can be recruited to the CNS during pupal development in response to glial immune activation and bacterial infection (Winkler et al., 2025). However, their involvement in CNS maintenance during aging remains largely unexplored. Whether partial defects in glial phagocytosis are sufficient to engage hemocyte recruitment to the brain has not been directly tested. Given that glial *drpr*-deficiency induces hyperactivation of immune signaling and inflammation (Elguero et al., 2023; Etchegaray et al., 2016), these observations raise the possibility that hyperactivation of glial immune signaling may promote recruitment of peripheral macrophages to the adult CNS.

Using a genetically encoded, membrane-tethered, pH-sensitive fluorescent reporter to visualize engulfed material within acidified phagolysosomal compartments, we find that *drpr*-deficient glia retain residual engulfment capacity but fail to efficiently degrade internalized material. As a result, enlarged, persistent phagocytic cargo accumulates with age. Impaired glial clearance also alters immune dynamics at the CNS periphery: peripheral hemocytes are increasingly recruited to the brain surface despite an intact BBB and engage in compensatory phagocytosis of debris.

Together, these findings reveal that age-associated declines in intracellular clearance efficiency, rather than complete phagocytic failure, are sufficient to reshape neuroimmune interactions and reveal a dynamic interface between CNS-resident glia and peripheral immune cells that emerges when homeostatic clearance declines.

## Results

We previously demonstrated that loss of *drpr* compromises apoptotic cell clearance, resulting in the accumulation of neuronal and glial corpses in the adult *Drosophila* brain as detected by TUNEL staining (Etchegaray et al., 2016). Notably, the number of TUNEL-positive cells remains relatively constant with age, suggesting that apoptotic cell death and clearance might reach a dynamic equilibrium rather than complete failure. This observation implies that residual phagocytic activity persists in *drpr*-deficient glia. However, whether such residual engulfment indeed persists had not been directly tested.

To directly assess ongoing engulfment and degradation, we employed pHRed, a genetically encoded, single-protein red fluorescent pH sensor that fluoresces selectively in acidic compartments (pH < 5.5), including phagosomes and lysosomes (Tantama et al., 2011). A membrane-tethered version of pHRed, generated by fusion to a CAAX motif, enables visualization of engulfed, membrane-bound material following acidification (Fig. 1A) (Hancock et al., 1992; Mondragon et al., 2019). We expressed membrane-tethered pHRed (hereafter, pHRed), selectively in neurons or glia to monitor the engulfment and processing of apoptotic materials in controls and flies with glial *drpr*-deficiency. Because glia are not the sole phagocytic population associated with the nervous system, we also considered whether peripheral immune cells (hemocytes) contribute to corpse clearance.

**Figure 1.**
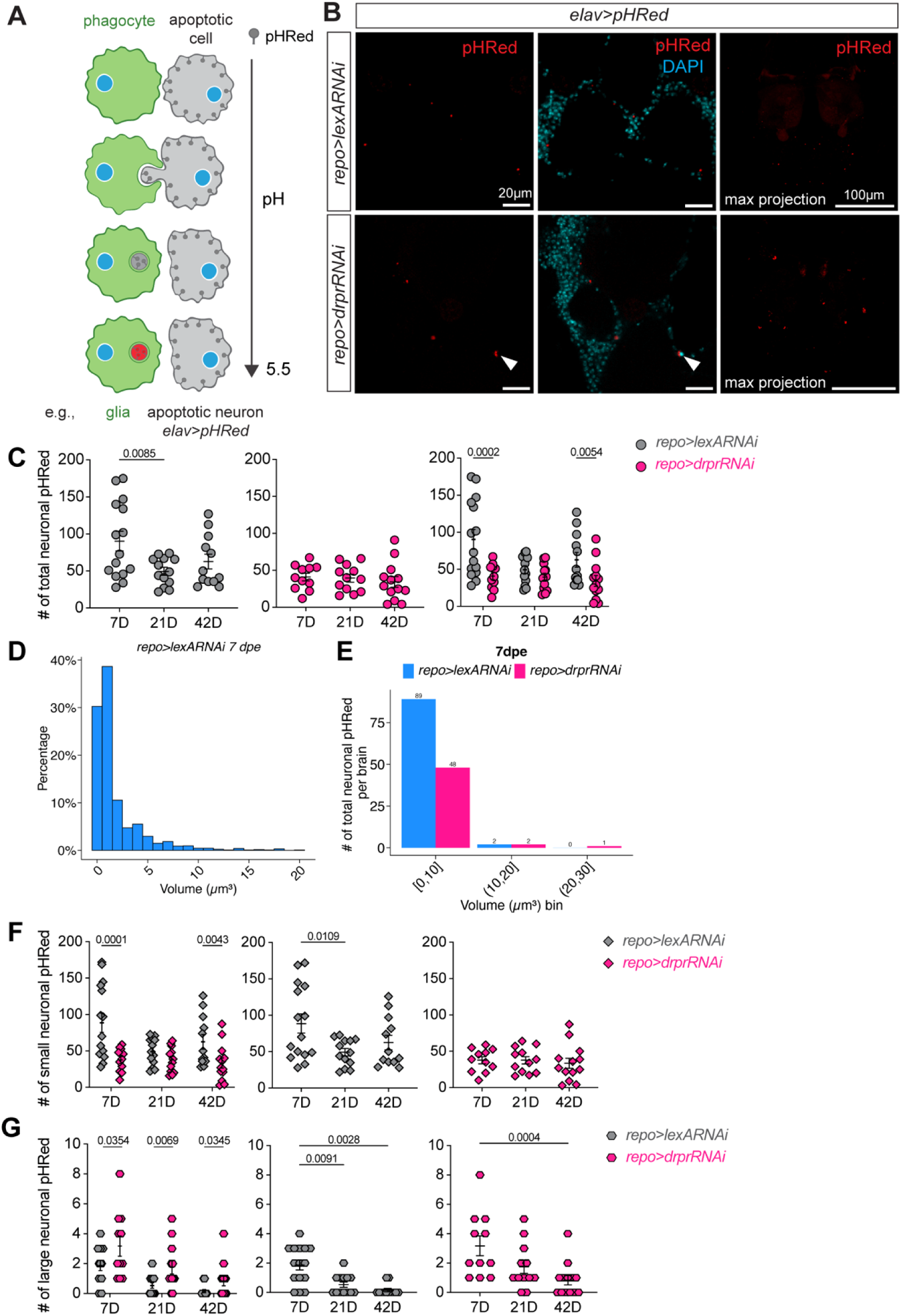
Neuronal pHRed dynamics in control and *drpr*-deficient brains across aging. (**A**) Diagram of pHRed as an acidification detector, adapted from Mondragon et al (Mondragon et al., 2019). (**B**) Immunofluorescence images of neuronal pHRed (red) and nuclei (DAPI, cyan) in 7 days post-eclosion (dpe) *repo>lexARNAi* and *repo>drprRNAi* brains. Scale bar, 20 µm. Arrowhead indicates pyknotic nucleus. (**C**) Quantification of average neuronal pHRed puncta per brain at 7, 21 and 42 dpe in *repo>lexARNAi* flies (7D, n=15; 21D, n=13; 42D, n=12) and *repo>drprRNAi* flies (7D, n=11; 21D, n=12; 42D, n=13). (**D**) Histogram showing the percentage distribution of neuronal pHRed volume in 7 dpe *repo>lexARNAi* brains (bin width = 1 µm^3^) (n=11). (**E**) Histogram of neuronal pHRed counts per brain in each 10 µm^3^ bin for *repo>lexARNAi* (blue) and *repo>drprRNAi* (red) brains. (**F**) Quantification of small (volume < 10 µm^3^) neuronal pHRed puncta in *repo>lexARNAi* and *repo>drprRNAi* brains across 7, 21 and 42 dpe. (**G**) Quantification of large (volume ≥ 10 µm^3^) neuronal pHRed puncta in *repo>lexARNAi* and *repo>drprRNAi* brains across 7, 21 and 42 dpe. Data information: Data represent mean ± SEM. Dots indicate numbers of pHRed puncta in individual brains. *P* values were determined by negative binomial regression.

### Neuronal engulfment is reduced and inefficient in *drpr*-deficient brains

To quantify neuronal engulfment, pHRed was expressed pan-neuronally using the *lexAop/lexA* system. Reporter specificity was validated by inducing traumatic brain injury (TBI), which triggers robust neuronal apoptosis. TBI resulted in a marked increase in neuronal pHRed fluorescence compared to sham controls, confirming that pHRed reliably reports acidified neuronal debris (Fig. S1A).

We next examined neuronal engulfment dynamics across lifespan in control and *drpr*-deficient brains. *drpr* dsRNA was driven in glia using the UAS/Gal4 system (*repo>drprRNAi)*, with *repo>lexARNAi* flies serving as controls. Brains were analyzed at 7, 21 and 42 days post-eclosion (dpe). Neuronal pHRed puncta localized predominantly to perinuclear regions, consistent with acidified cytoplasmic compartments (Fig. 1B).

In control brains, neuronal pHRed puncta varied with age, reaching a low point at 21 dpe (Fig. 1C). In contrast, *drpr*-deficient brains exhibited persistently low levels of neuronal pHRed puncta that did not significantly change with age. At both 7 and 42 dpe, *drpr*-deficient brains contained significantly fewer neuronal pHRed puncta than controls (Fig. 1C), indicating impaired engulfment and/or processing of neuronal material in the absence of functional *drpr*. The absence of a genotype difference at 21 dpe may reflect a transient balance between neuronal turnover and diminished glial clearance capacity, although the biological basis of this convergence remains unclear. Importantly, these data provide direct evidence that neuronal engulfment persists, albeit inefficiently, in *drpr*-deficient glia.

Beyond reduced abundance, neuronal pHRed puncta in *drpr*-deficient brains were conspicuously larger (Fig. 1B). In 7 dpe control brains, ∼98% of puncta were smaller than 10 µm^3^ (Fig. 1D). Based on this distribution, puncta were classified as “small” (< 10 µm^3^) or “large” (≥ 10 µm^3^). Volume distribution analysis revealed a pronounced enrichment of large puncta in *drpr*-deficient brains across all ages examined (Fig. 1E; Fig. S1, B-D). While small puncta followed trends similar to total counts (Fig. 1F), large neuronal pHRed puncta were significantly increased in *drpr*-deficient brains at all ages, despite declining with age in both genotypes (Fig. 1G). The accumulation of large, acidified neuronal remnants suggests defective phagolysosomal maturation and/or delayed degradation when *drpr* signaling is compromised (Etchegaray et al., 2016).

### Glial engulfment remains active but inefficient in *drpr*-deficient brains

Previous work showed that *drpr*-deficient glia themselves undergo apoptosis and are cleared over time, as indicated by colocalization of glial His2A-RFP with TUNEL signals and a decline in dying glia between 1 dpe and 10 dpe. To directly assess glial engulfment, we expressed pHRed in glia and quantified acidified structures in control and *drpr*-deficient brains at 3, 21 and 42 dpe. Glial pHRed signals exhibited marked heterogeneity in size, ranging from smaller puncta to entire glia fluorescing red (Fig. 2A). Total glial pHRed counts did not differ significantly between genotypes (Fig. 2B), although both showed age-dependent increases, indicating sustained phagocytic demand even in the absences of *drpr* (Fig. 2B).

**Figure 2.**
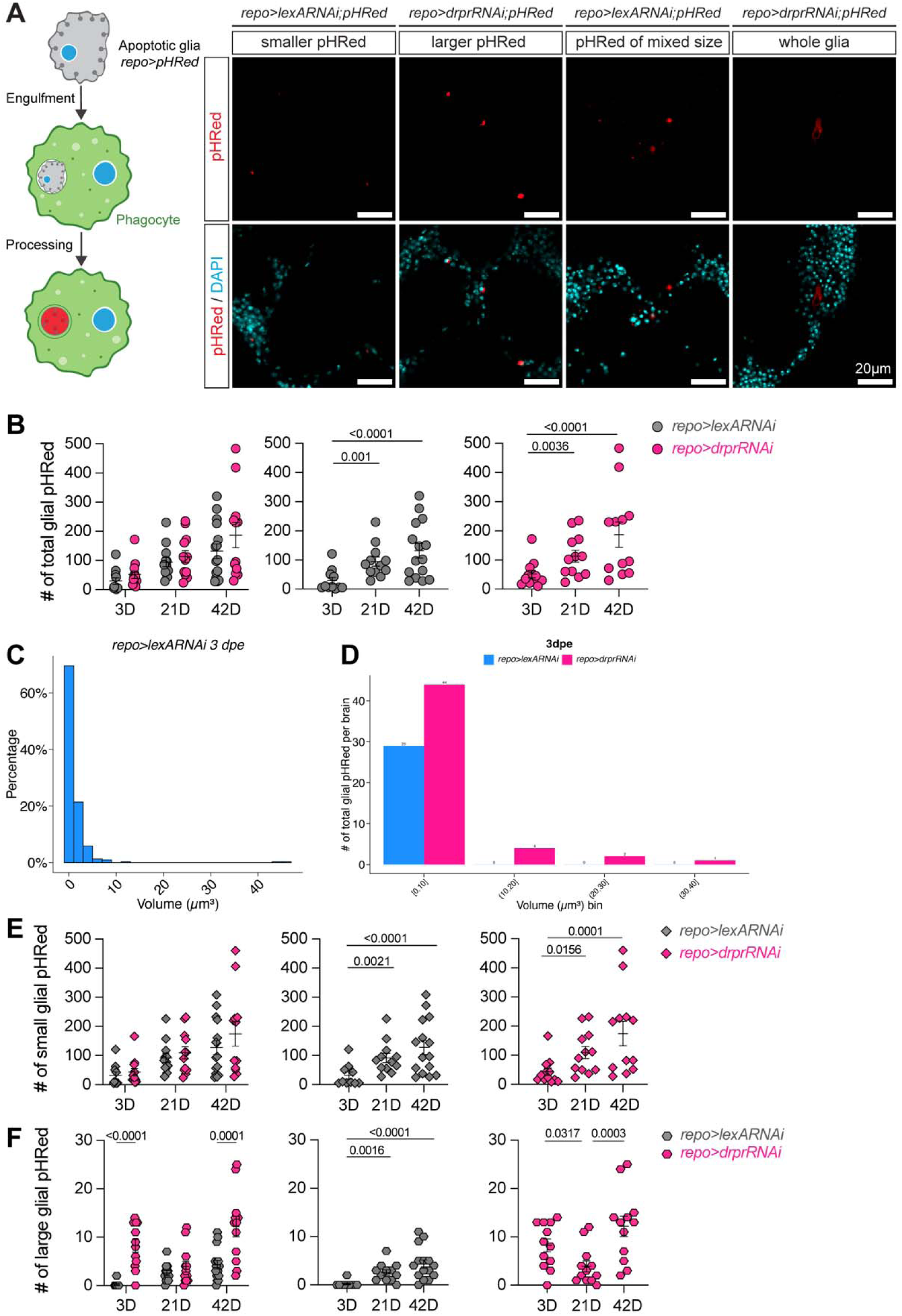
Glial pHRed dynamics in control and *drpr*-deficient brains across aging. (**A**) Diagram showing pHRed is expressed in glia. Representative immunofluorescence images of glial pHRed puncta of heterogenous size (red) and nuclei (cyan). Scale bar, 20 µm. (**B**) Quantification of average glial pHRed puncta per brain at 3, 21 and 42 dpe in *repo>lexARNAi* (3D, n=11; 21D, n=12; 42D, n=15) and *repo>drprRNAi* flies (3D, n=12; 21D, n=12; 42D, n=12). (**C**) Histogram showing the probability distribution of glial pHRed volume in 3 dpe *repo>lexARNAi* brains (bin width = 2 µm^3^; n=11). (**D**) Histogram of glial pHRed counts per brain in each 10 µm^3^ volume bin for *repo>lexARNAi* (blue) and *repo>drprRNAi* (red) brains. (**E**) Quantification of small (volume < 10 µm^3^) glial pHRed puncta in *repo>lexARNAi* and *repo>drprRNAi* brains across 3, 21 and 42 dpe. (**F**) Quantification of large (volume ≥ 10 µm^3^) glial pHRed puncta in *repo>lexARNAi* and *repo>drprRNAi* brains across 3, 21 and 42 dpe. Data information: Data represent mean ± SEM. *P* values were determined by negative binomial regression.

In 3 dpe control brains, 99.1% of glia pHRed puncta were smaller than 10 µm^3^ (Fig. 2C). *drpr*-deficient brains displayed a disproportionate increase in very large puncta (> 50 µm^3^) relative to controls (Fig. 2D; Fig. S2A-D). When stratified by size, small puncta largely mirrored total counts (Fig. 2E), whereas large puncta were significantly more abundant in *drpr*-deficient brains at 3 and 42 dpe (Fig. 2F). These findings indicate that although engulfment and acidification are initiated, degradation is delayed or incomplete in the absence of *drpr*.

Notably, the temporal dynamics of large puncta differed between genotypes. In controls, large puncta accumulated progressively with age, whereas in *drpr*-deficient brains they declined from 3 to 21 dpe and re-emerged again at 42 dpe (Fig. 2F), suggesting disrupted regulation of clearance capacity over time. This transient reduction may reflect partial compensatory mechanisms or turnover of dysfunctional glia, followed by age-associated exacerbation of phagocytic defects.

Large puncta located adjacent to pyknotic nuclei can often be attributed to single glial cells, whereas multiple small puncta clustered in a niche may represent repeated acidification events within glial cytoplasm. Because these small puncta are difficult to assign to individual glia, their raw counts are challenging to interpret. Thus, we also analyzed their relative abundance. *drpr*-deficient brains exhibited a lower proportion of small puncta at 3 and 42 dpe, with opposite age-related trends compared to controls (Fig. S2E). On the contrary, *drpr*-deficient brains exhibited a higher proportion of large puncta at 3 and 42 dpe compared to controls (Fig. S2F).

Notably, total glial pHRed counts exceeded neuronal pHRed counts in both controls and *drpr*-deficient brains (Fig 1C & 2B). Although neurons greatly outnumber glia in the adult brain, pHRed reports acidified intracellular compartments rather than cell number. We therefore hypothesize that the higher abundance of glial pHRed puncta reflects the intrinsic capacity of glia, as professional phagocytes, to generate multiple acidified compartments during the processing of diverse cellular material, including apoptotic cells (Poon et al., 2014), synaptic debris (Choi & Chung, 2025), axonal fragments (Watts et al., 2004) and damaged cellular components (Choi & Chung, 2025). In contrast, neuronal pHRed signal is expected to arise primarily following engulfment events and may be further constrained by lower phagocytic activity. Because pHRed is expressed intracellularly, glial pHRed puncta may also represent acidified endocytic or lysosomal compartments independent of phagocytosis. Nonetheless, we propose that the marked asymmetry between glial and neuronal pHRed abundance is consistent with a model in which glial phagocytic and degradative activity contributes disproportionately to the pool of acidified compartments detected by pHRed in the adult brain.

Taken together, whole-glia pHRed labeling likely reflects apoptotic or necrotic glia, whereas large puncta represent accumulated, undegraded material. These data demonstrate that *drpr*-deficient glia retain engulfment and acidification capacity but are inefficient at completing degradation. To distinguish partial from complete *drpr* loss, we quantified glial pHRed signals in *drpr*-null (*drpr*^Δ*5*^) flies (Fig. S2G). Compared to *drpr* knockdown, young *drpr*^Δ*5*^ brains exhibited fewer small puncta, while aged *drpr*^Δ*5*^ brains accumulated significantly more large puncta (Fig. S2G), indicating that complete loss of *drpr* exacerbates defects in glial degradation capacity.

### Glia continue to engulf neuronal and glial corpses in *drpr*-deficient brains

To confirm the identity of the phagocytic cells containing the pHRed^+^ vesicles, we expressed either cytoplasmic enhanced green fluorescent protein (EGFP) or membrane-tethered myristoylated GFP (myrGFP) in glia and assessed colocalization with neuronal or glial pHRed signals. In *drpr*-deficient brains, neuronal pHRed puncta colocalized with glial myrGFP (Fig. 3A&B), and glial pHRed puncta colocalized with glial EGFP (Fig. 3C&D). These results demonstrate that glia remain the principal phagocytes, engulfing both neuronal and glial corpses despite loss of *drpr*.

**Figure 3.**
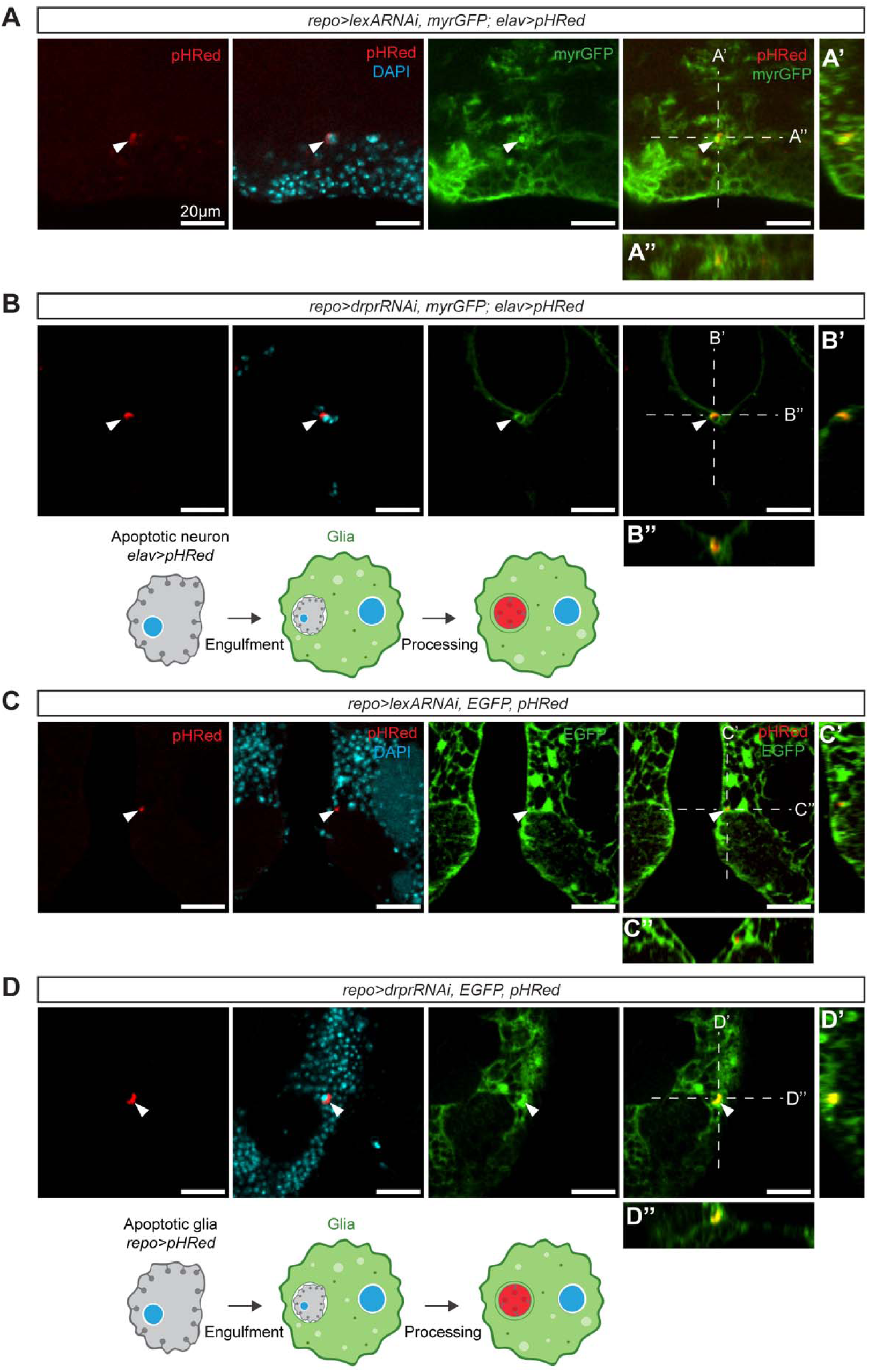
Glia engulf neuronal and glial corpses. (**A–B’’**) Immunofluorescence images and diagram of neuronal pHRed (red), nuclei (cyan), and pan-glial myristoylated GFP (myrGFP, green) in *repo>lexARNAi* (A) and *repo>drprRNAi* (B) brains. White arrowheads indicate colocalization of pHRed-positive neurons with myrGFP-labeled glia, with corresponding three-dimensional orthogonal view shown in (**A’, B’**) and (**A’’, B’’**). (**C-D’’**) Immunofluorescence images and diagram of glial pHRed (red), nuclei (cyan), and pan-glial cytoplasmic enhanced EGFP (green) *repo>drprRNAi* brain. White arrowheads indicate colocalization of pHRed-positive glia with EGFP-labeled glia, with corresponding three-dimensional orthogonal view shown in (C’, D’) and (C’’, D’’). Scale bar, 20 µm.

### Glial *drpr* deficiency promotes hemocyte adhesion to the CNS

In *Drosophila*, hemocytes have been shown to cross the BBB upon infection during the pupal stage, requiring immune deficiency (Imd)-dependent *pvf2* induction in glia and JAK/STAT dependent matrix metalloproteinases (MMP) activation in the BBB (Winkler et al., 2021, 2025). Because glial *drpr* deficiency induces corpse accumulation and upregulation of antimicrobial peptides downstream of Imd pathway (Elguero et al., 2023; Etchegaray et al., 2016), we next tested whether hemocytes were recruited to the adult brain. Hemocytes were labeled with anti-NimC1, and pan-glial EGFP was used to delineate the BBB. Analysis of whole-mount brains revealed hemocytes in three distinct locations: floating outside the brain, overlapping with the BBB, and rarely inside the CNS parenchyma (Fig. 4A). Internal hemocytes were observed randomly and no widespread hemocyte penetration into the brain parenchyma was observed (Fig. S3). Most hemocytes localized to the brain surface, often elongated and dorsally aligned, and persisted despite extensive washing, indicating stable adhesion (Fig. 4A). The number of external hemocytes may vary with dissection, therefore we quantified only those BBB-associated hemocytes in adult flies as a reliable indicator of brain-associated recruitment/adhesion.

**Figure 4.**
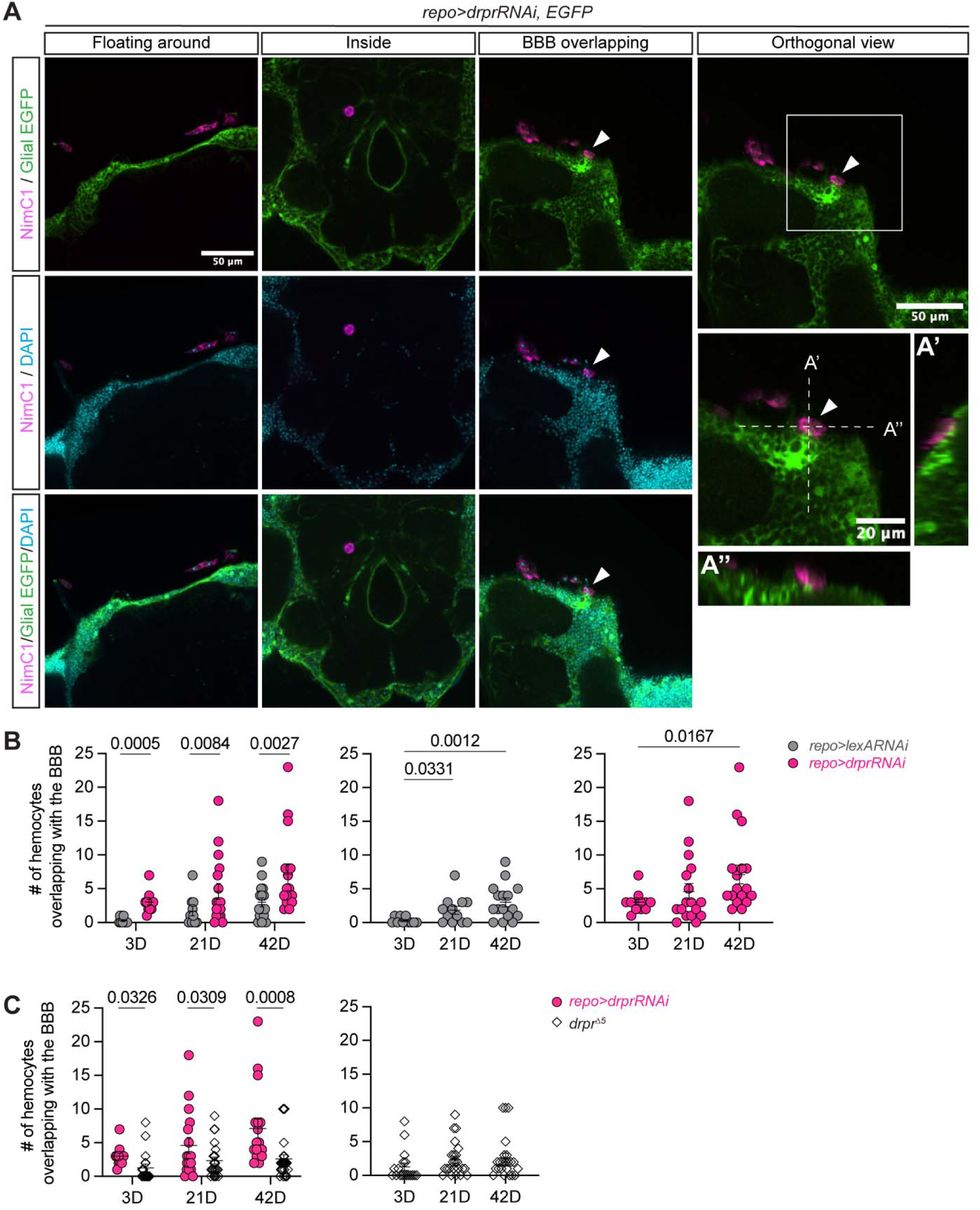
Hemocytes associate with the brain and the blood-brain barrier. (**A**) Immunofluorescence images showing hemocytes in three spatial contexts: floating outside the brain, within the brain parenchyma, and overlapping with the blood-brain barrier (BBB). Hemocytes stained with anti-NimC1 are shown in magenta, pan-glia in green and nuclei in cyan. White arrowheads indicate hemocytes contacting or overlapping with the BBB. Corresponding orthogonal views are shown in (**A**’) and (**A**’’). (**B**) Quantification of hemocytes overlapping with the BBB in *repo>lexARNAi* (3D, n=11; 21D, n=12; 42D, n=15) and *repo>drprRNAi* flies (3D, n=12; 21D, n=12; 42D, n=12) at 3, 21 and 42 dpe. (**C**) Comparison of the number of hemocytes overlapping with the BBB between *repo>drprRNAi* and *drpr*^Δ*5*^ flies (3D, n=18; 21D, n=23; 42D, n=24). Data information: Data represent mean ± SEM. Dots indicate individual brains. *P* values were determined by negative binomial regression. Scale bars are indicated on the figures.

*drpr*-deficient brains harbored significantly more BBB-associated hemocytes than controls at all ages examined (Fig. 4B). In both genotypes, BBB-associated hemocytes increased with age suggesting that aging and *drpr*-deficiency synergize to enhance hemocyte attraction (Fig. 4B). Importantly, deletion of *drpr* in hemocytes in a *drpr*^Δ*5*^ background significantly reduced BBB-associated hemocytes (Fig. 4C), demonstrating that hemocyte recruitment is likely partially dependent on *drpr* expression in hemocytes.

### *drpr*-deficiency elevates stress response in aged peripheral immune cells

Though glia retain partial phagocytic function in *drpr*-deficient brains, chronic corpse accumulation may overwhelm homeostatic capacity and influence peripheral immune physiology. In many pathological contexts, peripheral immune cells are often recruited to sites of pathology and they are accompanied by heightened cellular stress (Otxoa-de-Amezaga et al., 2019; Prinz & Priller, 2017; Yan et al., 2022; Zhang et al., 2019). To assess immune stress responses, we monitored oxidative stress using the GSTD-GFP reporter (Sykiotis & Bohmann, 2008).

In young adults, GSTD-GFP expression was minimal among hemocytes from both control and *drpr*-deficient brains. In contrast, aged *drpr*-deficient flies exhibited a pronounced increase in GSTD-GFP signal within hemocytes compared to age-matched controls (Fig. 5). These results indicate that glial *drpr* deficiency not only promotes hemocyte association with the CNS but also exacerbates oxidative stress in them during aging.

**Figure 5.**
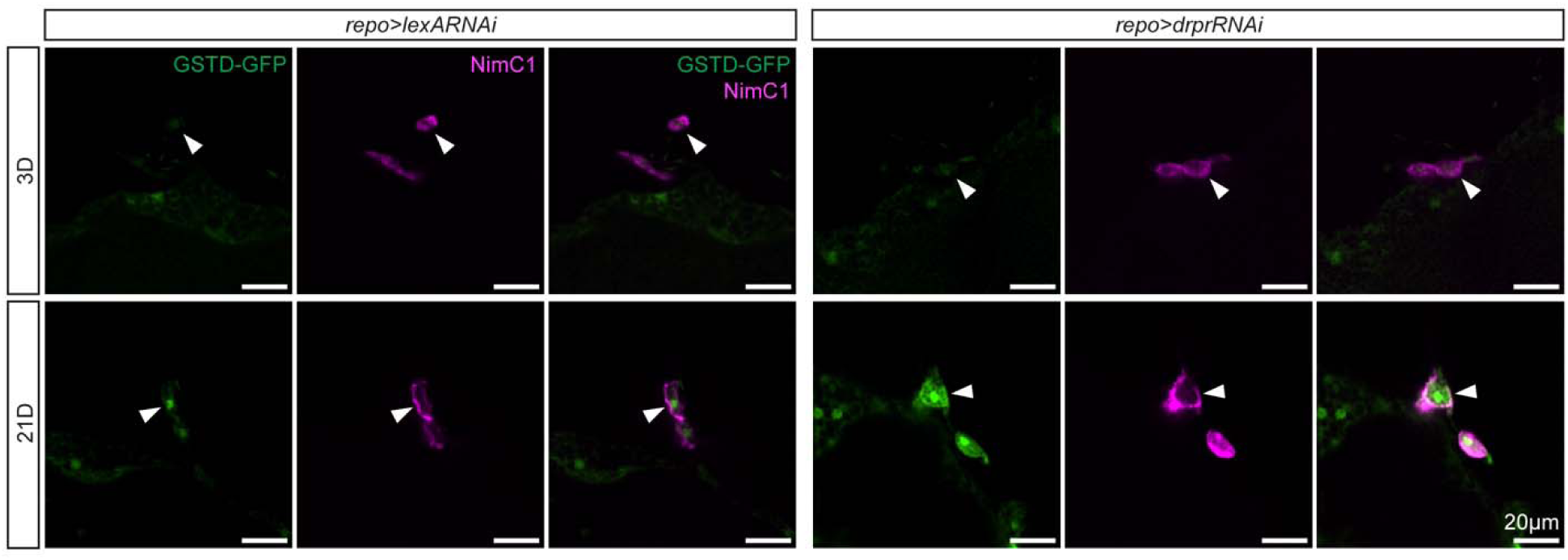
Oxidative stress in hemocytes. Immunofluorescence images showing oxidative stress reporter GSTD-GFP in hemocytes stained with anti-NimC1 (magenta) of *repo>lexARNAi* and *repo>drprRNAi* brains at 3 and 21 dpe.

### Peripheral hemocytes engulf glial material at the brain surface

Unexpectedly, some BBB-associated hemocytes contained glial pHRed puncta within their cytoplasm (Fig. 6A). Because hemocytes rarely enter the CNS parenchyma, these engulfed materials likely derive from border glia, including perineurial and subperineurial glia. The number of hemocytes containing glial pHRed increased with age and was consistently higher in *drpr*-deficient brains, particularly at 42 dpe (Fig. 6B).

**Figure 6.**
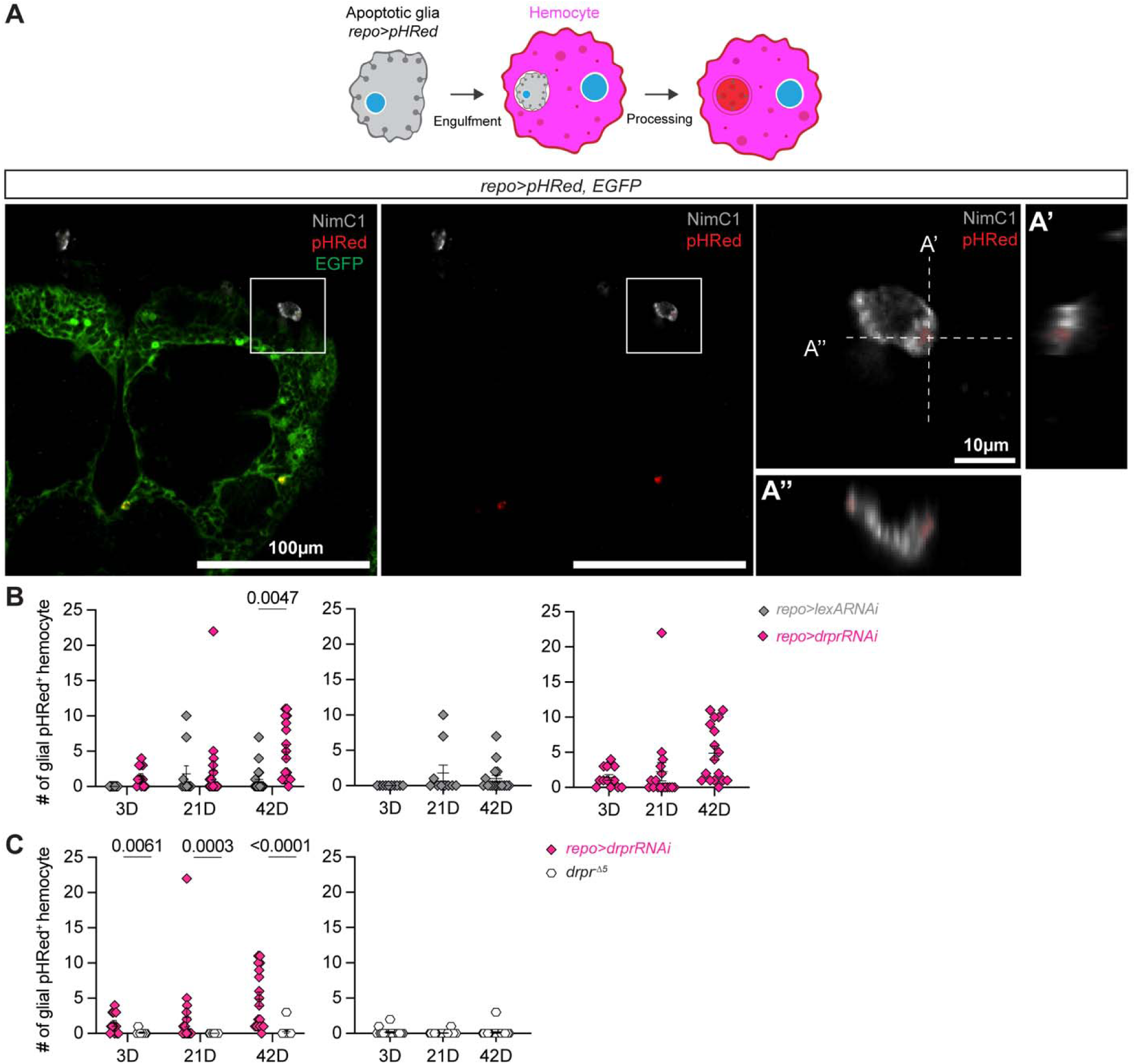
Hemocytes engulf glial corpses. (A) Representative immunofluorescence images and schematic illustrating hemocytes (grey) engulfing glial corpses expressing pHRed (red), with corresponding orthogonal view. (B) Quantification of hemocytes containing glial pHRed puncta in *repo>lexARNAi* (3D, n=11; 21D, n=12; 42D, n=15) and *repo>drprRNAi* (3D, n=12; 21D, n=12; 42D, n=12) brains at 3, 21 and 42 dpe. (C) Comparison of the number of hemocytes containing glial pHRed puncta between *repo>drprRNAi* and *drpr*^Δ*5*^ brains (3D, n=18; 21D, n=23; 42D, n=24). Data information: Data represent mean ± SEM. Dots indicate individual brains. *P* values were determined by zero-inflated negative binomial regression. Scale bars are indicated on the figures.

In control brains, hemocyte adhesion to the brain increased with age, but hemocyte-mediated engulfment of glial pHRed puncta remained rare (Fig. 6B). At early ages (e.g., 3 dpe), no hemocytes containing glial pHRed were detected. With aging, a small but noticeable increase in such events was observed, providing evidence that peripheral hemocytes can participate in phagocytosis in the aging brain. In *drpr*-deficient brains, significantly more hemocytes containing internalized glial pHRed puncta were detected compared to controls, particularly at 42 dpe (Fig. 6B). This phenomenon was abolished in *drpr*^Δ*5*^ brains lacking *drpr* in hemocytes (Fig. 6C), indicating hemocyte-mediated engulfment of glial material is *drpr*-dependent.

Collectively, these findings indicate that peripheral macrophages are attracted to, but do not invade, the CNS during aging, and that glial *drpr*-deficiency further amplifies this recruitment. Though hemocytes do not enter the brain parenchyma, they are nevertheless capable of phagocytosing glial material at the brain surface. How hemocytes access glia and mediate phagocytosis at this interface remains to be determined. Additionally, whether this hemocyte recruitment is ultimately beneficial or detrimental to CNS homeostasis requires further investigation.

### Blood-brain barrier integrity is preserved in *drpr*-deficient brains

The *Drosophila* BBB, formed by perineurial and subperineurial glia, maintain CNS homeostasis through septate junctions (Baumgartner et al., 1996). In adult *Drosophila*, BBB integrity is generally preserved, even under conditions that promote hemocyte association with the CNS (Winkler et al., 2021). Consistent with this, we observed no widespread penetration of hemocytes into the brain parenchyma (Fig. S3), suggesting that BBB function remains intact.

To directly determine whether *drpr* deficiency compromises BBB integrity, we assessed hemocyte localization, glial morphology, and dye permeability. Glial EGFP labeling revealed continuous and intact perineurial and subperineurial layers across all genotypes and ages examined (Fig. 7A). To evaluate barrier permeability, 10 kDa dextran was injected into the fly thorax, and its distribution relative to the neuropil was analyzed. Though dextran accumulated around the brain, no significant differences in dye penetration were observed between control and *drpr*-deficient brains at any age (Fig. 7B).

**Fig. 7.**
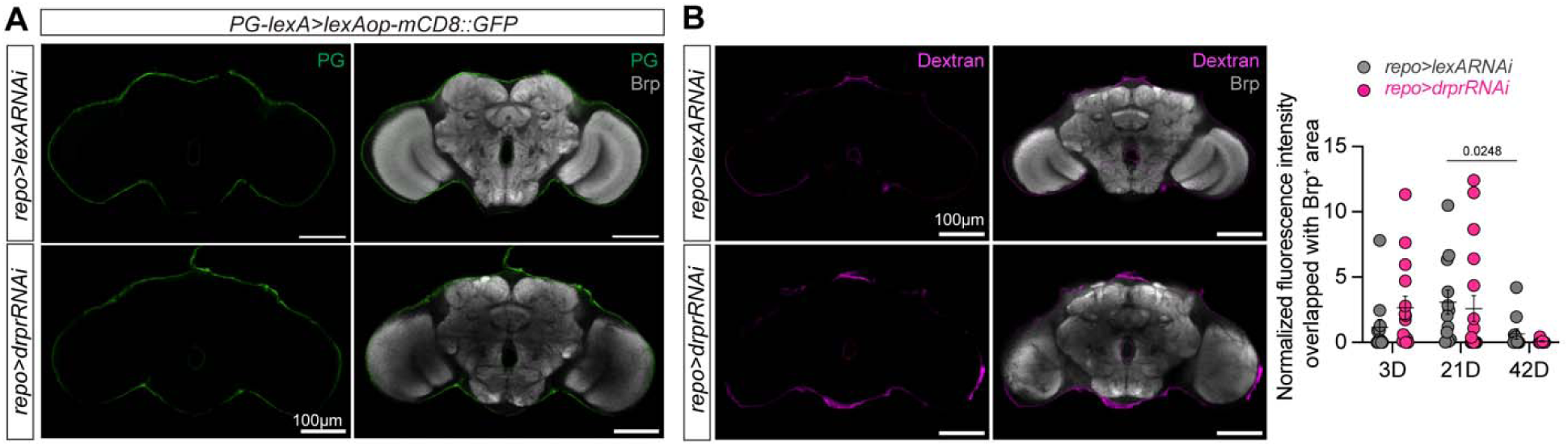
The blood-brain barrier remains intact upon *drpr* deficiency. (A) Visualization of the blood-brain barrier (BBB) in *repo>lexARNAi* and *repo>drprRNAi* brains. The BBB is demarcated by GFP expression in perineurial glia (PG) (*GMR85G01-lexA>lexAop-mCD8::GFP*, green). Brain neuropil structure is visualized with Bruchpilot (Brp, grey). (B) Assessment of BBB integrity by dextran penetration (magenta) into the brain (Brp, grey). Quantification shows dextran fluorescence intensity overlapping with the Brp-positive area, normalized to Brp area, in *repo>lexARNAi* (3D, n=12; 21D, n=12; 42D, n=11) and *repo>drprRNAi* flies (3D, n=15; 21D, n=18; 42D, n=9) at 3, 21 and 42 dpe. Scale bar, 100µm. *P* values were determined by one-way anova with Dunn’s multiple comparisons test.

Thus, the increased presence of hemocytes at the brain surface in *drpr*-deficient flies is not attributable to BBB breakdown, but instead likely reflects chemotactic attraction to inflammatory or degenerative signals emanating from the CNS.

## Discussion

Phagocytic clearance of apoptotic cells is a fundamental mechanism for maintaining tissue homeostasis, particularly in the nervous system where cell turnover, injury, and remodeling occur throughout life (Buss et al., 2006; Liu et al., 2022; Marín-Teva et al., 2012; Márquez-Ropero et al., 2020; Poon et al., 2014). In the adult *Drosophila* brain, glia act as phagocytes responsible for recognizing, engulfing, and degrading dying cells (Elguero et al., 2023; Etchegaray et al., 2016; Hilu-Dadia & Kurant, 2020; MacDonald et al., 2006; Purice et al., 2016). Loss of the engulfment receptor Draper in the null allele *drpr*^Δ*5*^ disrupts phagosome–lysosome fusion and impairs phagosomal acidification and maturation in both hemocytes and glia (Etchegaray et al., 2016; Petrignani et al., 2021; Shklover et al., 2015), indicating a role in phagocytic processing rather than cargo recognition alone. Consistent with this, corpses persist in glial *drpr*-deficient brain due to defective phagosome maturation, rather than failed engulfment, and enhancing lysosomal or metabolic pathways, such as JNK, TORC1 activation or Atg1 inhibition, can restore clearance capacity (Etchegaray et al., 2016; Petrignani et al., 2021). Our findings extend this framework by demonstrating that phagocytic dysfunction does not necessarily manifest as an all-or-none failure, but instead can arise as a progressive inefficiency in corpse processing, which in turn reshapes neuroimmune interactions during aging.

Incomplete clearance of cellular debris is increasingly recognized as a driver of inflammation across tissues. In the brain, persistent phagocytic cargo likely imposes a chronic metabolic and inflammatory burden on glia, altering their signaling state and interactions with other cell types (Safaiyan et al., 2016). Consistent with this idea, glial *drpr* deficiency is associated with age-dependent immune stress, reflected by increased oxidative stress signaling in peripheral hemocytes. Although hemocytes do not broadly infiltrate the CNS parenchyma, they are robustly recruited to the brain surface and adhere to the BBB in greater numbers when glial clearance is compromised. This recruitment is further amplified with aging, suggesting that glial phagocytic inefficiency synergizes with age-associated immune sensitization.

While our data support a link between impaired glial clearance and peripheral immune activation, the molecular signals mediating hemocyte recruitment remain unresolved. Multiple mechanisms likely contribute, including inflammatory cytokines (Winkler et al., 2021), extracellular matrix remodeling (Winkler et al., 2025), and stress-associated metabolites (Flowers et al., 2017). Dissecting these signals will be important for distinguishing direct consequences of defective phagocytosis from broader inflammatory changes associated with aging.

Although the brain has long been viewed as immunologically privileged, growing evidence highlights dynamic interactions at CNS interfaces (Smyth & Kipnis, 2025; Vara-Pérez & Movahedi, 2025). Our data show that peripheral hemocytes increasingly associate with the brain surface during aging and accumulate further when glial clearance is compromised. Chemotactic cues released by stressed or degenerating neurons and glia, potentially downstream of chronic immune pathway activation in *drpr*-deficient brains (Elguero et al., 2023), may drive this recruitment. The partial dependence of hemocyte recruitment on *drpr* function within hemocytes suggests that conserved engulfment machinery contributes to immune cell responsiveness to CNS-derived signals. At the same time, systemic immune aging may independently influence hemocyte behavior, acting in parallel with CNS-specific cues. Experimental strategies that decouple central and peripheral aging will be required to clarify their relative contributions.

Notably, hemocytes remain largely excluded from the CNS parenchyma yet can phagocytose glial material at the brain surface, likely involving perineurial and subperineurial glia. This behavior increases with age and glial *drpr* deficiency and requires *drpr* function in hemocytes. These findings support a model of barrier-associated “border clearance,” in which peripheral macrophages assist in removing damaged or dying glia at CNS interfaces without overt disruption of BBB integrity. Similar immune activities have been described for border-associated macrophages and perivascular immune cells in vertebrate systems, where they contribute to immune surveillance, debris clearance, and age-related inflammation (Drieu et al., 2022; Prinz & Priller, 2017; Schonhoff et al., 2023; Vara-Pérez & Movahedi, 2025; Zhan et al., 2025).

At the same time, our analysis does not rule out subtle or transient changes in BBB function below the resolution of the assays used. In addition, whether hemocyte-mediated clearance selectively targets compromised glia or certain glial subtypes or reflects opportunistic uptake of exposed material remains unclear. Determining the selectivity, timing, and functional consequences of border-associated phagocytosis will be essential for understanding its role in CNS maintenance.

Aging couples declining phagocytic efficiency with heightened immune reactivity (Deczkowska et al., 2018; Franceschi et al., 2017; Pluvinage & Wyss-Coray, 2020; Rawji et al., 2016; Safaiyan et al., 2016). Our findings highlight how these processes intersect: gradual impairments in corpse degradation generate a persistent source of debris that may amplify immune signaling over time. Rather than triggering acute pathology, this slow accumulation may promote chronic inflammation and increase vulnerability to neurodegenerative processes. Whether hemocyte-mediated border clearance ultimately alleviates glial burden or exacerbates inflammatory signaling likely depends on context, duration, and age.

Together, this study shows that inefficient corpse processing can persist alongside residual phagocytic activity, reshaping immune interactions at the CNS periphery during aging. By linking glial degradative capacity with peripheral immune engagement and barrier-associated phagocytosis, these findings provide a framework for understanding how failures in tissue maintenance influence neuroimmune homeostasis across the lifespan. They further suggest that strategies aimed at enhancing degradative efficiency, rather than solely promoting engulfment, may hold promise for mitigating age-associated neuroinflammatory states.

## Materials and Methods

### Fly strains and husbandry

All fly lines were maintained at 25°C on standard cornmeal-molasses food under a 12 h light/dark cycle. Aging flies were transferred to fresh food twice weekly to minimize microbial contamination and ensure consistent nutrition. The following fly strains were used: *UAS-lexA^RNAi^ (*BDSC #67947)) (Perkins et al., 2015), *UAS-drpr^RNAi^* (BDSC #67034) (MacDonald et al., 2006), *repo-Gal4/TM3* (BDSC #7415), *UAS-EGFP* (BDSC #5430), *UAS-myrGFP* (Pfeiffer et al., 2012), *UAS-pHRed* (Mondragon et al., 2019), *lexAop-pHRed* (generated in this study), *elav-lexA* (BDSC #94637), *drpr^Δ5^* (Freeman et al., 2003), *GMR85G01-lexA* (BDSC #54285), *lexAop-mCD8::GFP* (BDSC #32203). Experimental cohorts were aged to the specific time points indicated in the main text.

### Molecular cloning

pJFRC19-13XLexAop2-IVS-pHRed-CAAX (short for lexAop-pHRed) was made by replacing-myr::GFP in the plasmid pJFRC19-13XLexAop2-IVS-myr::GFP (Addgene: Plasmid #26224) with pHRed-CAAX (Table S1) and synthesized by GenScript (New Brunswick, NJ). The lexAop-pHRed was injected into embryos for PhiC31 transformation by BestGene (Chino Hills, CA).

### Dissection and histology

Flies were aged to designated days (3, 7, 21, 42 days post eclosion) and equal numbers of male and female flies were dissected. Flies were first fixed with 4% paraformaldehyde (PFA) in phosphate-buffered saline with 0.3% Triton-X 100 (0.3% PBT) for 3 hours at RT and washed four times with 0.3% PBT within one hour. Brains were dissected in phosphate-buffered saline (PBS) at room temperature (RT). Brains were permeabilized with 0.3% PBT for one hour at RT or overnight at 4°C. Samples were blocked in PBANG (0.3% PBT, 0.5% bovine serum albumin (BSA), and 5% normal goat serum (NGS)) for 1 hour at RT or overnight at 4°C and then incubated in primary antibody diluted in PBANG for 2 nights at 4°C. Samples were then washed in 0.3% PBT four times for a total of one hour at RT, before being incubated in secondary antibody diluted in PBANG for 2 nights at 4°C in the dark. Samples were washed with PBT twice and 1X PBS twice for a total of one hour at RT, followed with being transferred to Vectashield mounting media with DAPI and stored in the dark at 4°C until mounting on slides. All the incubation and washes were done with rotation. The following primary antibodies were used: rabbit anti-GFP (Torrey Pines, 1:100), mouse anti-NimC1 (gift from István Andó, 1:100; now DSHB, Dm N1/9) (Kurucz et al., 2007). The following secondary antibodies were used: goat anti-rabbit Alexa Fluor 488 (JacksonImmuno #,111-545-003 1:200), goat anti-mouse Alexa Fluor 647 (JacksonImmuno #115-605-003, 1:200).

### Traumatic brain injury (TBI)

Traumatic brain injury (TBI) was induced as previously described (Dixon et al., 2025). Briefly, 7-day-old adult flies were placed in an empty vial with a 1-inch vertical clearance. A foam plug was inserted at the 1-inch mark from the bottom of the vial, and the vial was mounted onto the high-impact trauma (HIT) device. Injuries were administered at a 90° angle, with each experimental group receiving three strikes. Flies were allowed to recover for 1 day before brain dissection.

### Blood-brain barrier integrity assay

Flies at 3, 21, 42 dpe were anesthetized on a CO_2_ pad and injected using a microinjector (Tritech Research, Inc,) equipped with 1-mm needle. Injections were performed under a dissecting microscope with either 1x PBS or 10k Da Dextran conjugated with Alexa Fluor™ 647 (Thermo Fisher Scientific). An average volume of 100 ± 25 nl was injected into the lateral thorax between the wing socket and the haltere. The fluorescent dextran was diluted in 1x PBS to a final concentration of 10 mg/ml. After injection, flies were allowed to recover in standard food vials at 25 °C overnight (∼16 hours). Whole flies were fixed in 0.3% PBT containing 4% PFA and processed for dissection and histology as described above. Neuropils were stained mouse anti-nc82/Brp (DSHB, 1:100).

Brains were imaged using a Nikon AX confocal microscope from the anterior side, acquiring 51 slices at a 2 µm step size with a 20x objective and 1024×1024 pixel resolution. Images were analyzed in Fiji/ImageJ using a custom macro. Colocalization of permeabilized dextran with Brp staining was quantified, and normalized dextran pixel intensity was calculated by dividing the total colocalized dextran signal by the Brp-positive area.

### Microscopy

Brains mounted on glass slides were imaged using a Nikon AX confocal microscope. Images were acquired with a 40× objective lens (NA 0.95; working distance, 0.21 mm). For each brain, 61 optical sections were collected at a resolution of 1024 × 1024 pixels.

### Statistical analysis

The number of pHRed-positive puncta was quantified using a custom Fiji (ImageJ) macro. Statistical analyses were performed using Poisson or negative binomial regression models in R (v4.4.2), as appropriate for the data distribution. Statistical analyses were graphed and analyzed in either Graphpad Prism (v9) or R as indicated in corresponding figure legend.

## Data and materials availability

All data are available in the main text or the supplementary materials.

## Acknowledgments

We thank István Andó for sharing anti-NimC1 antibody and Ranjan Mithal for technical support.

## Author contributions

GL and KM conceived the study, designed the experiments, and interpreted the results. GL, IA and CYS performed the experiments. GL wrote the paper and KM reviewed and edited the paper. KM acquired funding. GL performed data curation, data analysis, and visualization.

