## Supplemental figures for "Phagocytosis deficient glia display phagosome processing defects and macrophage recruitment to the brain of adult *Drosophila melanogaster*"

Guangmei Liu *et al.*

Kimberly McCall,

**This PDF file includes:**

Figs. S1 to S3
Tables S1

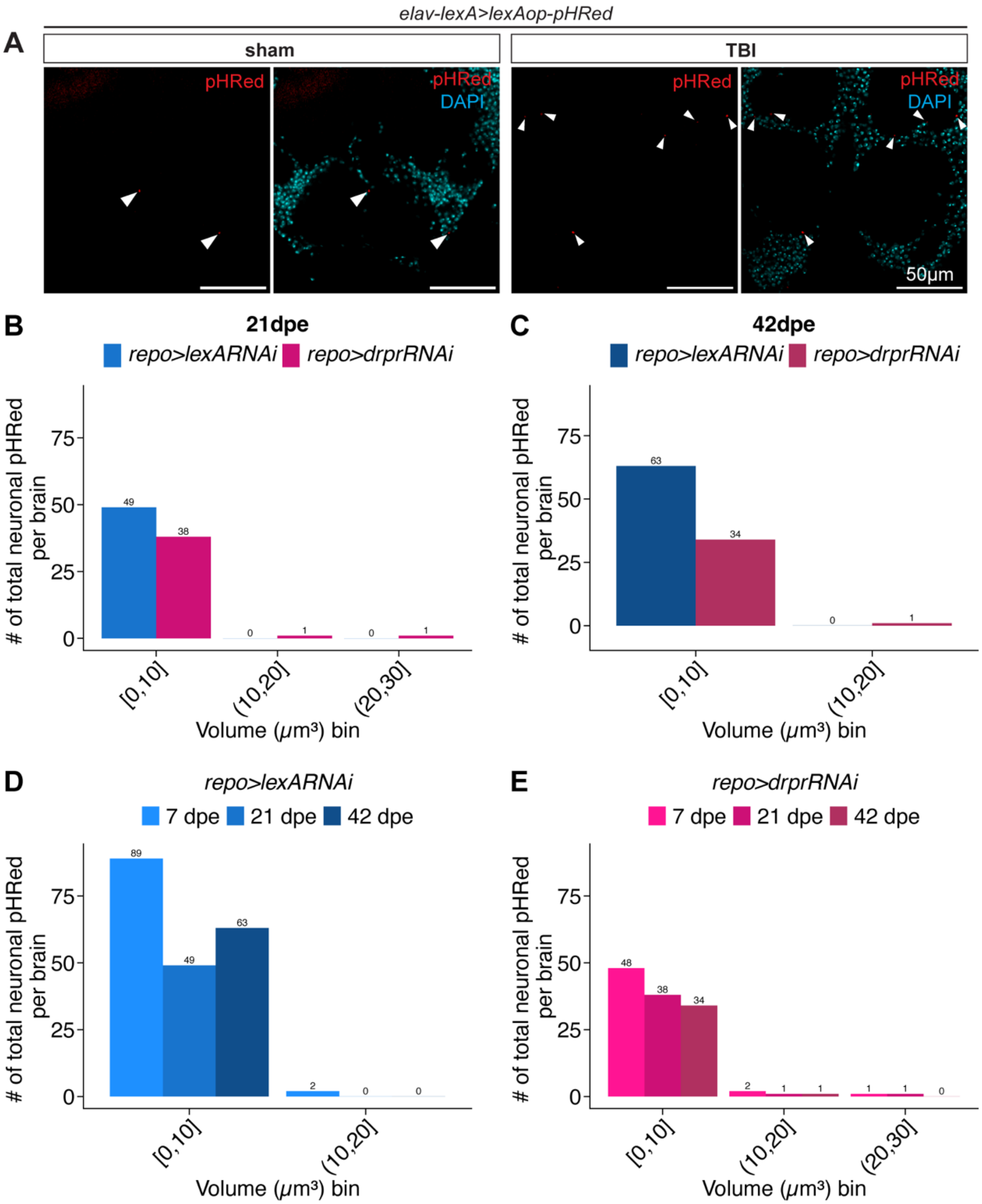

elav-lexA>lexAop-pHRed
A
sham
TBI
pHRed
DAPI
50μm
B
21dpe
repo>lexARNAi
repo>drprRNAi
### of total neuronal pHRed per brain
Volume (μm³) bin
[0,10]
(10,20]
(20,30]
49
38
0
1
0
1
C
42dpe
repo>lexARNAi
repo>drprRNAi
63
34
0
1
D
repo>lexARNAi
7 dpe
21 dpe
42 dpe
89
49
63
2
0
0
E
repo>drprRNAi
7 dpe
21 dpe
42 dpe
48
38
34
2
1
1
1
1
0

**Fig. S1. Neuronal pHRed size distribution.**

(**A**) Validation of *lexAop-pHRed* expression using the *elav-lexA* driver in flies subjected to traumatic brain injury (TBI) and sham controls. White arrowheads indicate pHRed puncta. Scale bar, 50 µm. (**B, C**) Histograms of neuronal pHRed counts per brain in each 10 $\mu m^3$ bin for *repo>lexARNAi* (blue) and *repo>drprRNAi* (red) brains at 21(**B**) and 42 (**C**) days post-eclosion (dpe). (**D, E**) Histogram of neuronal pHRed counts per brain in each 10 $\mu m^3$ bin for *repo>lexARNAi* (**D**) and *repo>drprRNAi* (**E**) brains across 7, 21 and 42 dpe.

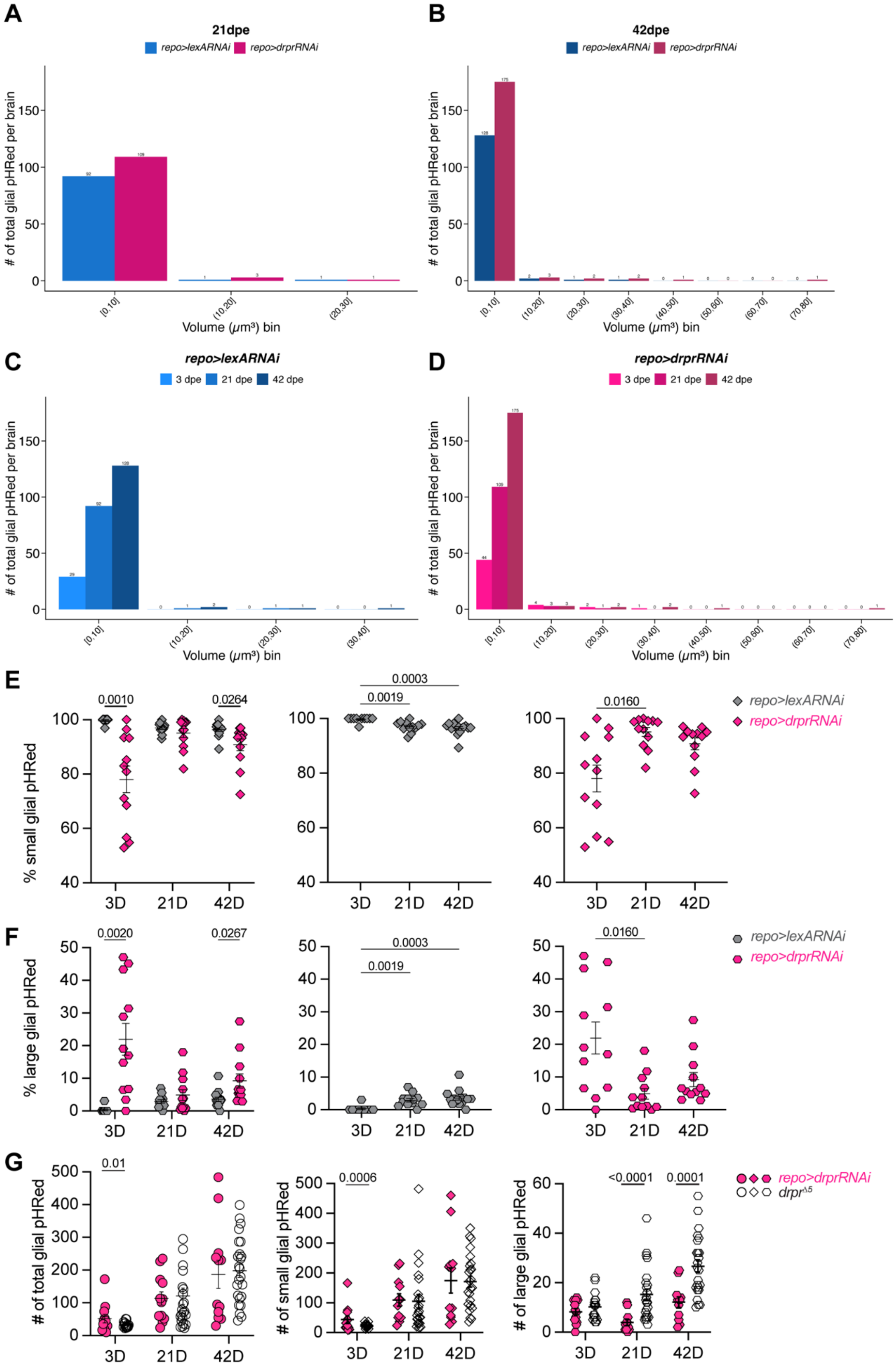
A
21dpe
repo>lexARNAi
repo>drprRNAi
### of total glial pHRed per brain
Volume (μm³) bin
B
42dpe
repo>lexARNAi
repo>drprRNAi
### of total glial pHRed per brain
Volume (μm³) bin
C
repo>lexARNAi
3 dpe
21 dpe
42 dpe
### of total glial pHRed per brain
Volume (μm³) bin
D
repo>drprRNAi
3 dpe
21 dpe
42 dpe
### of total glial pHRed per brain
Volume (μm³) bin
E
% small glial pHRed
0.0010
0.0264
0.0003
0.0019
0.0160
3D
21D
42D
repo>lexARNAi
repo>drprRNAi
F
% large glial pHRed
0.0020
0.0267
0.0003
0.0019
0.0160
3D
21D
42D
repo>lexARNAi
repo>drprRNAi
G
### of total glial pHRed
0.01
### of small glial pHRed
0.0006
### of large glial pHRed
<0.0001
0.0001
3D
21D
42D
repo>drprRNAi
drpr^Δ5

**Fig. S2. Glial pHRed size distribution.**

(**A, B**) Histograms of glial pHRed counts per brain in each 10 $\mu m^3$ volume bin for *repo>lexARNAi* (blue) and *repo>drprRNAi* (red) brains at 21(**A**) and 42 (**B**) dpe. (**C**) Histogram of glial pHRed counts per brain in each 10 $\mu m^3$ bin for *repo>lexARNAi* brains across 3, 21 and 42 dpe. (**D**) Histogram of glial pHRed counts per brain in each 10 $\mu m^3$ bin for *repo>drprRNAi* brains across 3, 21 and 42 dpe. (**E, F**) Percentage of small (**E**) and large (**F**) glial pHRed puncta in *repo>lexARNAi* and *repo>drprRNAi* brains across 3, 21 and 42 dpe. (**G**) Comparison of total, small and large number of glial pHRed puncta between *repo>drprRNAi* and $drpr^{-/-}$ flies (3D, n=18; 21D, n=23; 42D, n=24). Dots indicate individual brains. Data represent mean ± SEM. *P* values were determined by negative binomial regression.

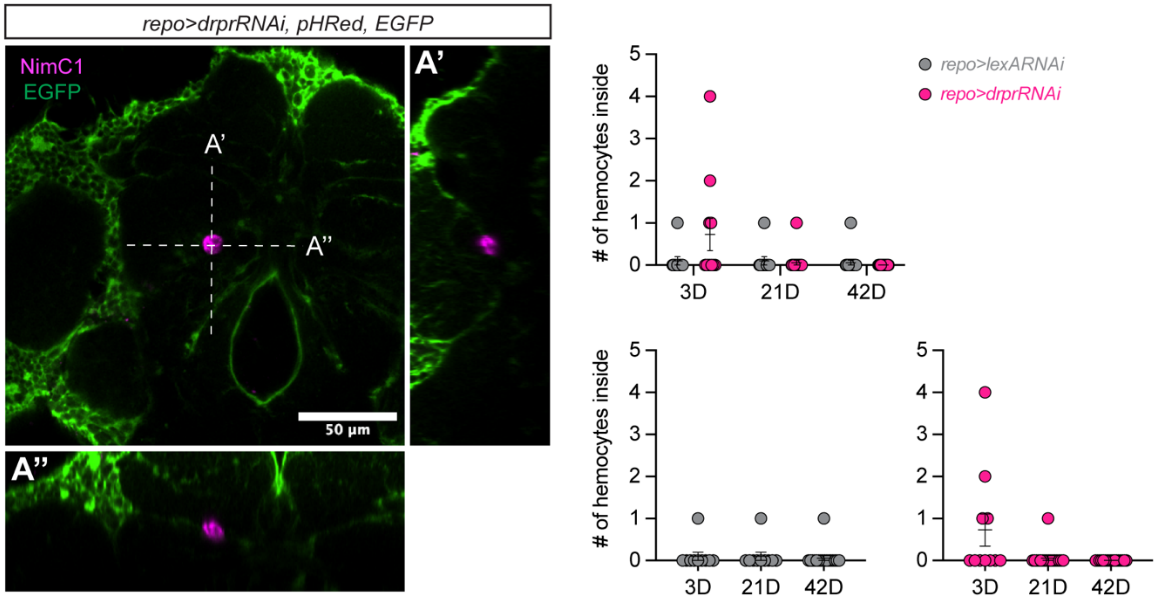


**Fig. S3. Few hemocytes are found inside the brain.**

(**A**) Orthogonal view of brains expressing EGFP in pan-glia (green) and labeling hemocytes (magenta). (**B**) Quantification of hemocytes located within the brain parenchyma in *repo>lexARNAi* (3D, n=11; 21D, n=12; 42D, n=15) and *repo>drprRNAi* brains (3D, n=12; 21D, n=12; 42D, n=12) at 3, 21 and 42 dpe. Data information: Data represent mean ± SEM. Dots indicate individual brains. *P* values were determined by zero-inflated negative binomial regression.

**Table S1. Construct and primer sequences of pHRed-CAAX.**

| **pHRed-CAAX (5’-3’)** |
| --- |
| ATGGTTTCCGTTATCGCAAAGCAAATGACCTATAAAGTTTATATGTCTGGCACCGT<br>TAACGGCCACTACTTCGAGGTTGAAGGTGACGGCAAGGGCAAGCCTTACGAGGGC<br>GAGCAGACCGTTAAGCTGACCGTTACCAAAGGCGGCCCACTGCCTTTCGCCTGGG<br>ATATCCTGTCCCCTCAGCTCCAATACGGCTCCATCCCTTTCACCAAATACCCCGAG<br>GACATCCCTGACTACTTTAAGCAGAGCTTCCCAGAGGGCTACACCTGGGAGCGCT<br>CTATGAACTTCGAGGACGGCGCAGTTTGTACCGTTTCCAACGACAGCTCCATCCAG<br>GGTAACTGCTTCATCTACAACGTTAAGATTTCCGGCGAGAACTTCCCACCTAACGG<br>CCCAGTTATGCAGAAGAAAACCCAAGGCTGGGAGCCCAGCACCGAACGCCTGTTT<br>GCCCGCGACGGTATGCTGATTGGTAACGATTATATGGCCCTGAAACTGGAAGGCG<br>GCGGTCACTACCTGTGCGAGTTCAAGAGCACTTACAAGGCTAAAAAGCCTGTTCG<br>CATGCCTGGCCGCCACGAAATCGACCGCAAGCTGGATGTTACCTCTCACAACCGC<br>GACTACACCAGCGTTGAGCAGTGTGAAATCTCTATCGCCCGCCACTCTCTGCTGGG<br>GACAGGCAACTCCGCCGACGGTGGTGGCTCTGGTGGTGCTAGCAAAGAAAAGATG<br>AGCAAAGATGGTAAAAAGAAGAAAAAGAAGTCAAAGACAAAGTGTGTAATTATG<br>TAA |
| **pHRed-CAAX forward primer:** |
| CACCATGGTTTCCGTTATCGCAAAG |
| **pHRed-CAAX reverse primer:** |
| TTACATAATTACACACTTTGTCTTTGAC |
